# Intratumoral Heterogeneity of Vimentin Modulates Nuclear Mechanotransduction, DNA Damage Response and Cancer Cell Survival

**DOI:** 10.1101/2025.06.27.661506

**Authors:** Infante Elvira, Terriac Emmanuel, Gelin Matthieu, Siegfried Hugo, Pereira David, Roca Vanessa, Varet Hugo, Khalilian Sara, Rietveld Reinier, Asnacios Atef, van Bodegraven Emma J, Etienne-Manneville Sandrine

## Abstract

Vimentin, a major intermediate filament protein, is essential for maintaining cellular integrity and regulating cytoskeletal dynamics. Its upregulation is a hallmark of epithelial-to-mesenchymal transition (EMT), a process that enhances cancer cell migration, invasion, and metastatic potential. However, single-cell transcriptomic analyses of glioblastoma, the most common and aggressive primary brain tumor, reveal that vimentin expression exhibits significant intratumoral heterogeneity, reflecting diverse cellular subpopulations that may contribute to tumor plasticity, therapy resistance, and disease progression. Here, we show that the absence of vimentin alters nuclear mechanotransduction in response to compression, leading to chromatin remodelling and profound changes in gene expression in cancer cells. Remarkably, we demonstrate that external compressive forces, akin to vimentin deficiency, disrupt DNA damage response pathways. This impairment compromises DNA damage sensing and repair, bypassing DNA damage checkpoints and apoptosis. Consequently, vimentin-negative tumor cells exhibit increased survival in response to physical stress and DNA damage, potentially driving radioresistance and further amplifying intratumoral heterogeneity.

## Introduction

Cancer progression from tumour formation to dissemination involves a complex interplay of genetic, biochemical, and mechanical changes. While genetic and biochemical cues have long been recognized as key drivers, mechanical cues—such as increased tissue stiffness, compressive forces, and shear stresses—have emerged both as significant prognostic factors in cancer progression (Barnes et al., 2018; Martino et al., 2018) and as potential therapeutic targets (Kaukonen et al., 2016; Nia et al., 2020). The contributions of tumour mechanics, encompassing the mechanical properties of both the extracellular environment and tumour cells, are particularly relevant in cancer progression.

The mechanics of cells, including cancer cells, relies mainly on their cytoskeleton, composed of actin, microtubules and intermediate filaments. Actin filaments and microtubules are highly conserved cytoskeletal components present in all cell types, maintaining a relatively uniform composition across different tissues. In contrast, intermediate filaments, such as vimentin, display cell type-specific expression and dynamically adapt to environmental cues, allowing cells to modulate their structural and functional properties in response to physiological and pathological conditions (Etienne Manneville, 2018). Notably, intermediate filament composition can change during processes like cell differentiation or cancer progression. Shifts in protein expression often serve as markers for cellular identity and pathological states and have been linked to patient prognosis (Sharma et al., 2019). The best known example is Vimentin IFs, a major cytoskeletal component of motile mesenchymal cells and a hallmark of epithelial-to-mesenchymal transition (EMT) found in metastatic tumours of epithelial origin (Danielsson et al., 2018; Guo et al., 2012). Beyond serving as biomarkers, IFs play a key role in cellular and tissue mechanics, influencing properties like stiffness and resilience (Etienne-Manneville, 2018; Infante and Etienne-Manneville, 2022; van Bodegraven and Etienne-Manneville, 2021). Changes in IF composition are associated with cell invasive capacity and tumour spreading (Leduc and Etienne-Manneville, 2015), highlighting their key role in the interplay between cell mechanical properties and extracellular mechanical cues. Vimentin expression have been shown to promote cell migration, invasion, and altered mechanics in numerous cancers, including glioblastoma (GBM) (Mendez et al., 2010; Patteson et al., 2019; Sivagurunathan et al., 2022; Strouhalova et al., 2020; Thievessen et al., 2015; van Bodegraven et al., 2023), however, the significance of a potential low-vimentin reservoir within the tumour has never been investigated.

GBMs are the most aggressive and devastating primary brain tumours, with limited treatment options and poor prognoses. As GBM grows in the brain enclosed within a hard skull, compression of the brain tissue and of the tumour mass is a typical feature and a key cause of the clinical symptoms observed in patients with brain cancer. Accumulating evidence from *in vitro* studies, animal models, and patient tissues demonstrate a clear link between the mechanical properties of tumour tissue and GBM aggressiveness (Barnes et al., 2018; Miroshnikova et al., 2016), as well as increased therapeutic resistance (Erickson et al., 2018). More recently, compression forces were shown to promote GBM progression. The mechanical properties of glioma cells also evolve during tumour progression (Alibert et al., 2020; Alibert et al., 2021), suggesting changes in their cytoskeletal composition. It also highlights the need to better understand the dynamic interplay between mechanics of the cells and their microenvironment, as this relationship has profound implications for tumour progression and therapy resistance.

In this study, we demonstrate the coexistence of both vimentin-positive and vimentin-negative cells within GBM samples and show that while vimentin expression was shown to facilitate GBM cell invasion, its absence may paradoxically enhance cell resistance to stress. These findings underscore the unexpected dualistic role of vimentin in GBM, suggesting it may serve as a valuable molecular marker for distinguishing tumour cell subpopulations with distinct behaviors. Understanding this heterogeneity is critical for developing more precise and effective therapeutic strategies tailored to the complexity of GBM.

## Results

### Heterogenous vimentin expression in glioblastoma

The GBM genetic landscape has been shown to evolve dynamically over space and time, producing an extraordinary degree of cellular complexity and heterogeneity within individual tumours (Greenwald et al., 2024; Qazi et al., 2017; Reinartz et al., 2017; Sottoriva et al., 2013). To determine whether vimentin expression in GBM also showed intratumoral heterogeneity, we analysed published patient data from single-cell RNA sequencing (scRNAseq) from a large integrated GBM scRNAseq dataset of Isocitrate dehydrogenase (IDH)-wild-type GBM (Ruiz-Moreno, 2022). This analysis showed a heterogenous expression of vimentin in all tumour samples (**Fig. 1A**) and variations in the percentage of vimentin-positive cells across patients (**Supplementary Figure Fig.1A**). Differential gene expression analysis of cells from the tumour core versus the tumour periphery (Darmanis et al., 2017) revealed that vimentin-negative cells are significantly more abundant in the core than in the periphery (**Fig. 1B**). This finding suggests that both vimentin-positive and –negative cells persist in high-grade tumour and that low or absence of vimentin expression may be more advantageous to cells within the tumour core.

**Figure 1.**
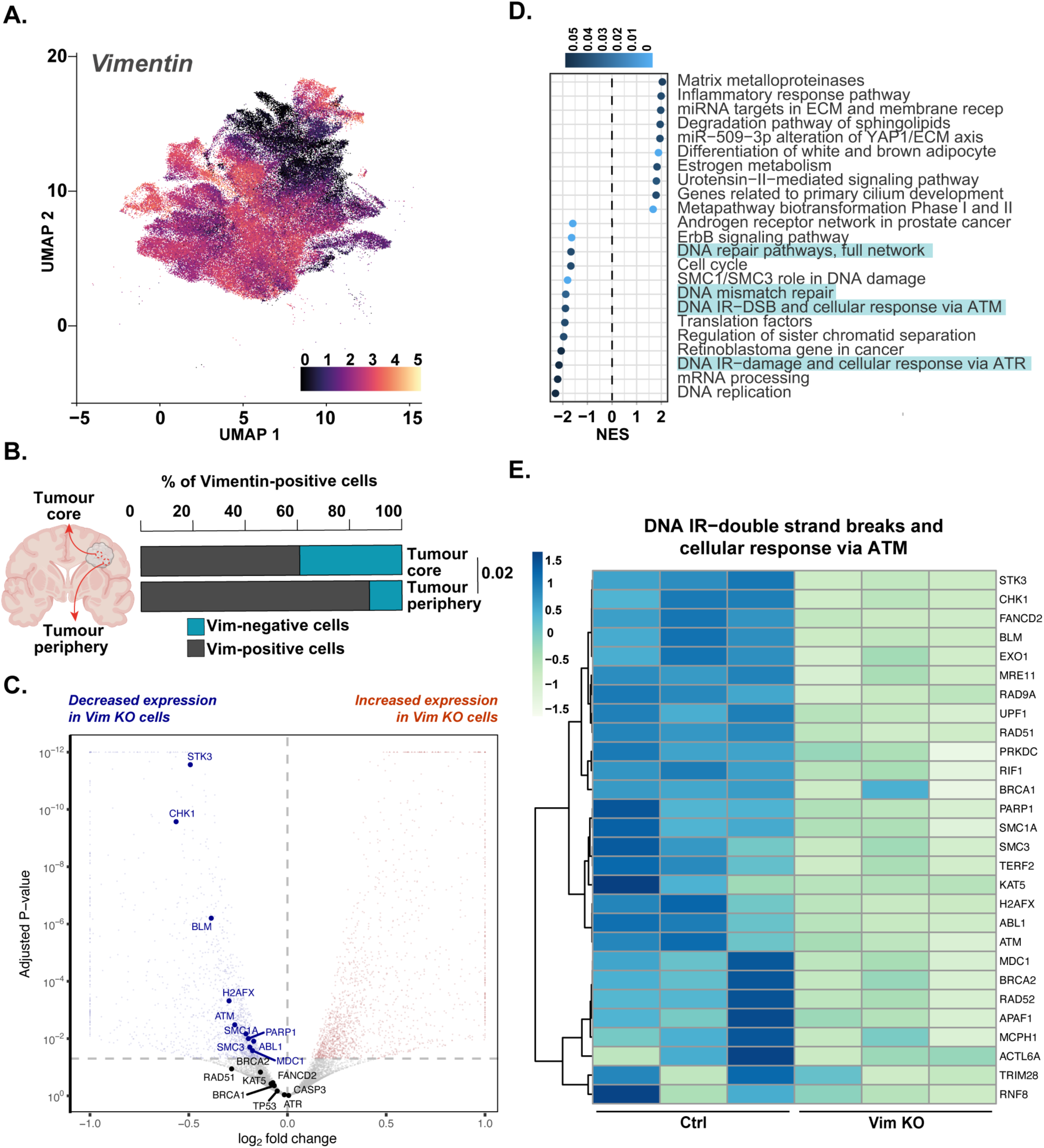
Vimentin heterogeneity in GBM. **A.** Uniform Manifold Analysis Approximation and Projection (UMAP) analysis shows the different neoplastic cell populations present within a large integrated GBM single-cell RNA sequencing dataset. A fire LUT was applied to highlight vimentin heterogenous expression within these cell populations, with low expression denoted in black and high expression in light yellow. **B.** Percentage of vimentin-positive neoplastic cells in the tumour core and tumour periphery. Vimentin-positive and -negative neoplastic cell percentages of four (P1-4) different GBM grade IV IDHwt patients for which location information is shown (Darmanis et al., 2017). Data were analysed by chi-square test of independence. **C.** Volcano plot of the differential expression analysis between control and vimentin-depleted cells. Differentially expressed (DE) genes were selected based on adjusted *p*-value of ≤ 0.05. The volcano plot is restricted to log2FC between -1 and +1 and adj P-values >10^(-12)^. Upregulated genes (red dots) and downregulated genes (blue dots) are shown. **D.** Pathway enrichment analysis based on WikiPathways gene-sets using the GSEA method. NES stands for Normalised Enrichment Score. **E.** Heatmaps providing a visual representation of gene expression of the DNA IR-double strand breaks and cellular response via ATM pathways in control versus Vim KO U251-MG cells. The heatmap is based on the variance-stabilised transformed count matrix and rows have been re-orderd thanks to a hierarchical clustering using the correlation distance and the Ward aggregation criterion. Gene names are reported. Colour scale ranges from -1.5 to +1.5 as the rows of the matrix have been centered. High expression is denoted in dark blue, while low expression in light green.

### Loss of vimentin impairs of DNA damage repair (DDR) pathways in GBM cells

To study whether the absence of vimentin modifies specific cell characteristics, we compared the transcriptome of U251-MG GBM cells expressing high vimentin levels and a vimentin knockout (Vim KO) U251-MG cell population obtained using a CRISPR/Cas9 approach (van Bodegraven EJ et al., 2023) (**Supplementary Figure Fig.1B)**. Transcriptomic analysis revealed a strong modification of gene expression profiles of Vim KO cells compared to control cells (1646 downregulated genes (p_adj_<0.05) and 1877 upregulated genes (p_adj_<0.05)) (**Fig. 1C**). Interestingly, among the most down regulated genes, we found genes belonging to the Gene Ontology (GO) terms related to DNA repair pathways, including DNA repair pathways, full network, DNA mismatch repair, DNA IR (ionizing radiation-induced)-DSB (double strand breaks) and cellular response via ATM (Ataxia telangiectasia mutated) as well as DNA IR−damage and cellular response via ATR (ATM and Rad3 related) (**Fig. 1D**). A more detailed analysis of the identified genes showed a particularly striking downregulation of genes encoding critical sensors and regulators of the DNA repair machinery in Vim KO cells (**Fig. 1E** and **Supplementary Figure Fig. 1C**). Notably, this includes genes encoding 1) the H2A histone family member X, H2AX, a key histone variant involved in the detection and signalling of double-strand breaks (DSBs), 2) ATM, a critical kinase which initiates signalling cascades that regulate DSB repair and replication stress response, and 3) the checkpoint kinase 1, Chk1, a central kinase in the DNA damage response (DDR) which plays an essential role in cell cycle arrest. Additionally, several *RAD* genes, including *RAD9A*, which can be phosphorylated by ATM in response to DNA damage, and controls G_1_/S checkpoint(Yin et al., 2004) were also downregulated. A significant downregulation of ABL1, a gene with DNA-binding activity also implicated in DNA damage responses and apoptosis, together with a downregulation of SMC3 and SMC1A involved in DNA repair, was observed in vimentin deficient cells. The analysis also identified reduced expression of PARP1, a key enzyme in base excision repair (BER) and an important player in the repair of single-strand breaks (SSBs). The downregulation of these genes suggests a substantial impairment of both homologous recombination and non-homologous end joining (NHEJ) pathways in vimentin-deficient cells.

To confirm the downregulation of major DDR proteins, we performed western blot analysis in Vim KO U251-MG cells. Results showed a significant reduction of ATM, Chk1 and H2AX protein expression compared to control U251-MG cells (**Fig. 2A**). These observations were also confirmed by transiently downregulating vimentin expression using two different small interference RNA (siRNA) technology (**Fig. 2B**). We also validated our observation in patient-derived GBM cells (U3117), which express high levels of vimentin. In these cells, vimentin depletion by CRISPR/Cas9 also reduced the expression level of ATM, Chk1 and H2AX (**Supplementary Figure Fig. 2A**).

**Figure 2.**
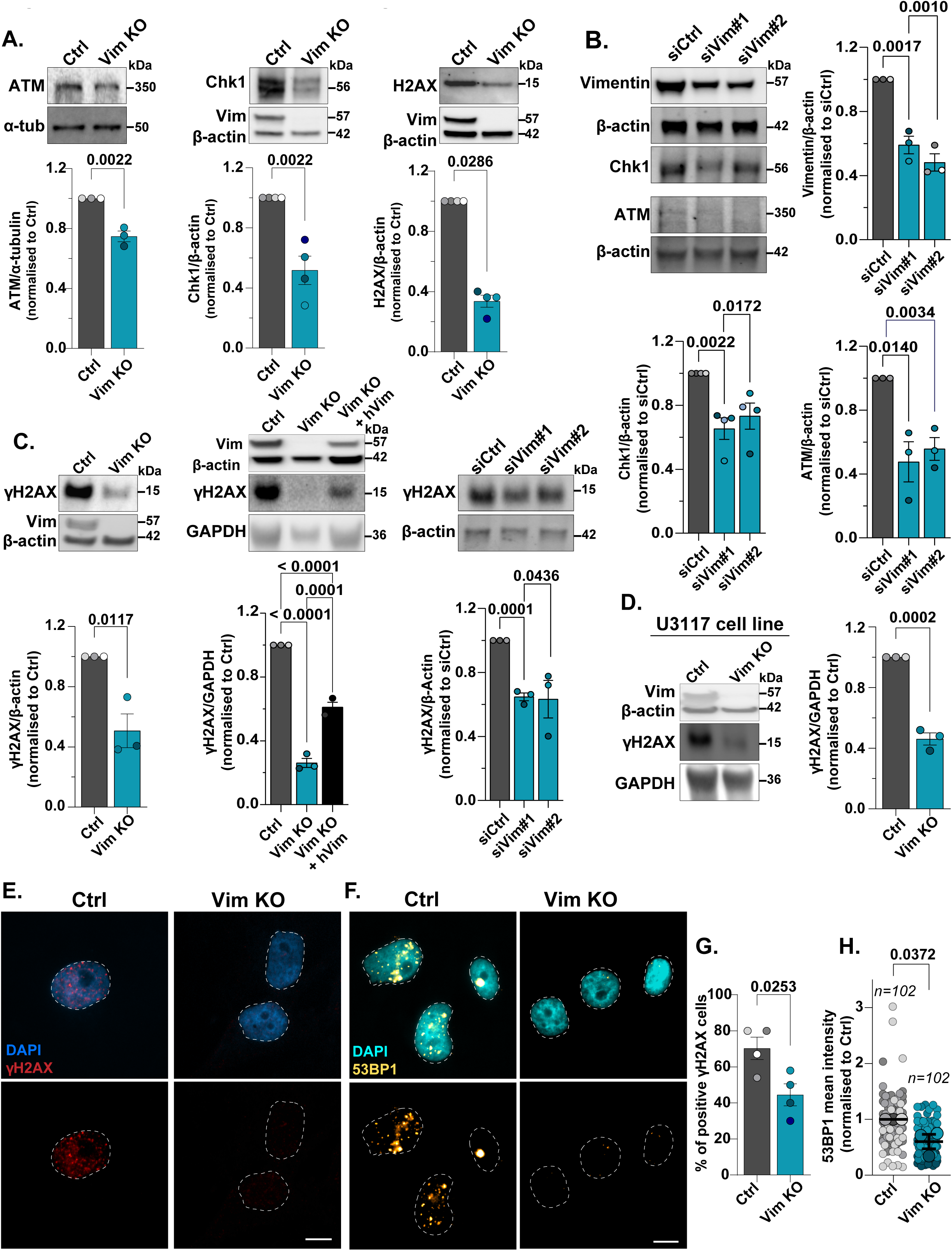
DDR proteins are downregulated in Vim KO cells. **A.** Expression of ATM, Chk1 and H2AX in control and vimentin knockout U251-MG cells was analysed by western blot and relative expression was quantified using Fiji. Data were acquired from three independent experiments (ATM), and from four independent experiments (Chk1 and H2AX) and analysed using a two-tailed *t*-test. **B.** U251-MG cells were transfected with siRNAs targeting vimentin. Levels of vimentin, Chk1 and ATM proteins were analysed by western blot and relative expression quantified using Fiji. Data were acquired from three independent experiments, and from four independent experiments (Chk1) and analysed using a one-way ANOVA test. **C.** Expression of γH2AX in control, Vim KO and vimentin-rescued cells and relative expression were quantified using Fiji. Data were acquired from three independent experiments and analysed by a two-tailed *t*-test (left graph) or a one-way ANOVA test. **D.** γH2AX and vimentin expression in control and Vim KO U3117 primary cells were analysed by western blot and relative expression quantified using Fiji. Data were acquired from three independent experiments and analysed by a two-tailed *t*-test. **E and F.** U251-MG cells depleted or not for vimentin were plated on glass coverslips and stained for (E) γH2AX or (F) 53BP1 and DAPI to visualise the nuclei (scale bar 10 µm). **(G)** The percentage of γH2AX positive cells was quantified. Data were acquired from four independent experiments and analysed by a two-tailed *t*-test. **(H)** SuperPlots of the mean intensity of 53BP1 staining. Single data points are represented as small dots in grey (Ctrl) or blue (Vim KO). The mean of each repeat is symbolized by bigger dots. Data were acquired from three independent experiments and analysed by a two-tailed *t*-test. *n*=total number of cells per condition.

DNA double strand breaks (DSB) are followed by the phosphorylation of H2AX by kinases such as ATM and ATR (Bonner et al., 2008). This newly phosphorylated protein, γH2AX, is the first step in recruiting and localizing DNA repair proteins. Western blots as well as immunofluorescence experiments showed a decrease in γH2AX in Vim KO U251-MG cells, in patient-derived GBM cells (U3117), and in U251-MG siRNA-mediated vimentin-depleted cells compared to control cells (**Fig. 2C-G**). This result was also confirmed in five different single clones of Vim KO U251-MG obtained using CRISPR/Cas9 (**Supplementary Figure Fig. 2C**). Moreover, re-expression of human vimentin in Vim KO U251-MG cells led to a partial but highly significant rescue of γH2AX (**Fig. 2C**).

Following activation of the DNA damage response (DDR) by ATM, Nuclear factor kappaB (NF-κB) is recruited to the nucleus leading to the transcription of genes involved in cell survival, inflammation, and repair (Miyamoto, 2011). We observed a decrease of nuclear NF-κB in vim KO cells compared to control cells (**Supplementary Figure Fig. 2E**). Finally, we also checked the expression level of the p53-binding protein 1, 53BP1, a crucial component of DNA DSB signalling and repair, which surprisingly was not included in the GO list of DNA repair pathway genes. Immunofluorescence analysis showed that 53BP1 was also strongly reduced in Vim KO cells, further confirming the general inhibition of DNA damage sensing and DDR pathways (**Fig. 2F and 2H**). Together, these results show a profound alteration of the DDR pathway in cells lacking vimentin, and suggest that vimentin-negative GBM cells are unable to detect and efficiently repair DNA damage.

### Loss of vimentin impairs of DNA damage repair (DDR) pathways in adenocarcinoma cells

To determine if vimentin expression also influenced DNA repair gene expression in the context of epithelial-to-mesenchymal transition, we used a breast adenocarcinoma cell line (MDA-MB-231). These cells lack epithelial markers like E-cadherin and instead express mesenchymal markers such as vimentin. Knockdown of vimentin using siRNA led to the downregulation of DNA damage response (DDR) proteins, including Chk1 and H2AX (**Supplementary Figure Fig. 2B**), consistent with observations in GBM cells. Finally, downregulation of γH2AX was also observed in vimentin-depleted MDA-MB-231 cells (**Supplementary Figure Fig. 2D**).

This demonstrates that vimentin’s impact on DDR pathway regulation is robust across diverse cellular contexts, including tumours of epithelial origin.

### Loss of vimentin impairs cell responses to damage damage

To determine if the DNA damage sensing machinery was functionally altered in vimentin-deficient cells, we artificially induced DNA damage by exposing control and Vim KO U251-MG cells to ultraviolet (UV) radiation, a well-established and potent inducer of DNA lesions. To directly assess DNA damage, we performed a comet assay under alkaline conditions, which detects both single-strand and double-strand breaks (Olive and Banath, 2006). UV treatment significantly increased the comet aspect ratio in control, Vim KO and vimentin-rescued cells with no significant differences between conditions (**Fig. 3A**), indicative of DNA break formation irrespectively of the presence or absence of vimentin. However, while in control U251-MG cells, γH2AX levels increased markedly following UV exposure, as expected, reflecting the activation of the DNA damage response, Vim KO cells exhibited a significantly attenuated increase in γH2AX, confirming impaired DNA damage sensing (**Fig. 3B**). Given this strong influence of vimentin expression on DNA damage sensing and DDR pathways, we next examined its impact on GBM cell survival following DNA damage. Western blot analysis of the proliferation marker PCNA revealed comparable baseline expression between non-treated control and Vim KO cells (**Fig. 3C**). In control cells PCNA levels significantly dropped following UV exposure. In contrast, in Vim KO cells UV exposure do not induce any statistically significant changes in PCNA levels (**Fig. 3C**).

**Figure 3.**
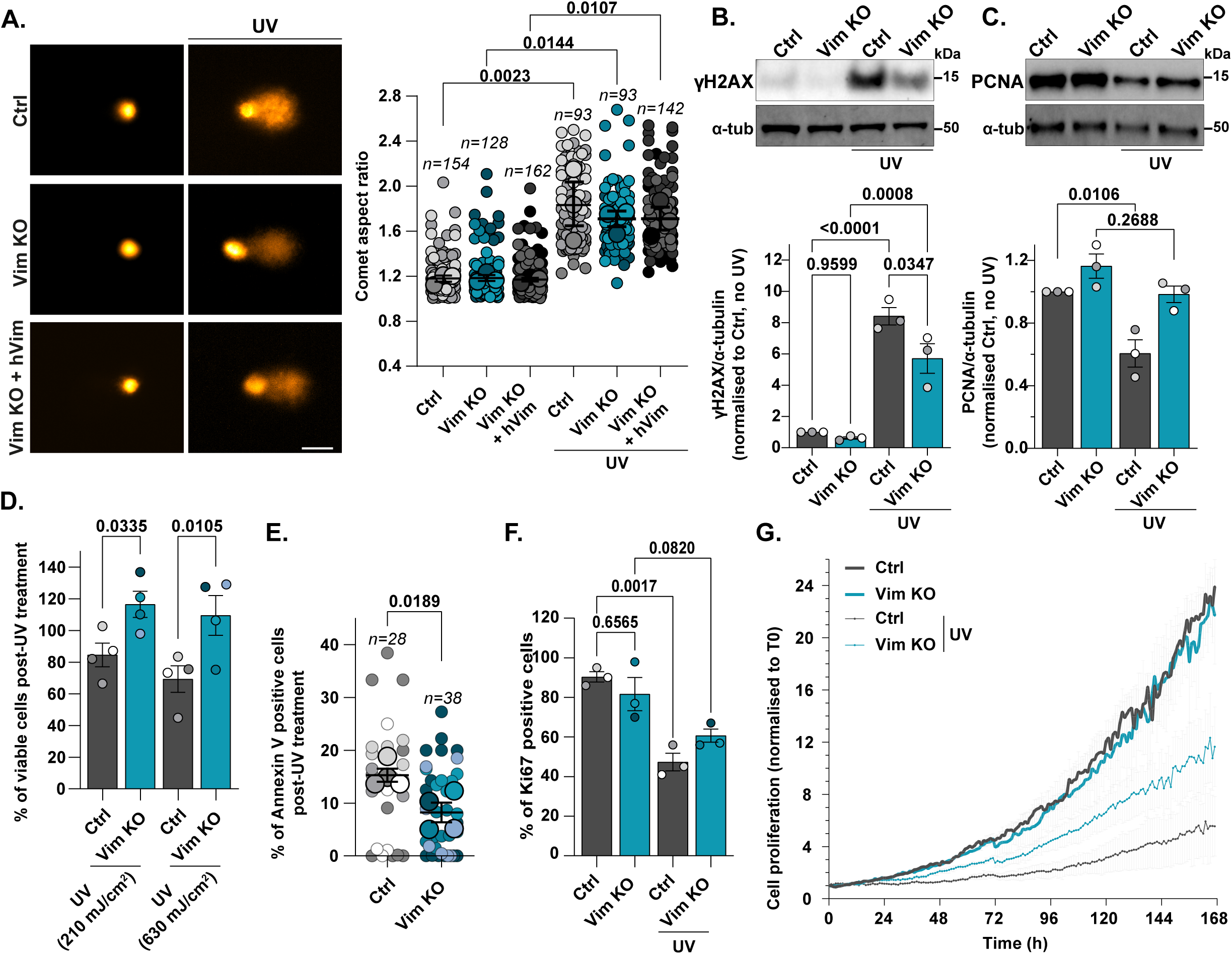
Vimentin-depleted cells are less sensitive to DNA damage. **A.** (Left) Representative images of U251 control, vimentin-depleted cells and vimentin-rescued cells irradiated or not with 1.05 J/cm^2^ UV light and after 3h subjected to alkaline comet assay (scale bar 20 µm). (Right) SuperPlots of the comet aspect ratio. Single data points are represented as small dots in grey (Ctrl), blue (Vim KO) or black (Vimentin rescue). The mean of each repeat is symbolized by bigger dots. Data were acquired from three independent experiments and analysed by a one-way ANOVA test. *n*=total number of cells per conditions. **B and C.** U251 cells depleted or not for vimentin were subjected to 210 mJ/cm^2^ UV light irradiation, then lysed after 24h and protein level of γH2AX and PCNA analysed by western blot and quantified using Fiji. Data were acquired from three independent experiments and analysed by a one-way ANOVA test. **D.** MTT assay of control and Vim KO cells performed 24h after 210 mJ/cm^2^ or 630 mJ/cm^2^ UV light irradiation. Data were acquired from four independent experiments and analysed by a one-way ANOVA test. **E.** Quantification of percentage of annexin V-positive cells in control and Vim KO U251 cells after 24h of UV light irradiation. SuperPlots of the % of Annexin V positive cells. Single data points are represented as small dots in grey (Ctrl) or blue (Vim KO). The mean of each repeat is symbolized by bigger dots. Data were acquired from three independent experiments and analysed by a two-tailed *t*-test. *n*=total number of cells per condition. **F.** Quantification of ki67 positive cells 24h after 210 mJ/cm^2^ UV light irradiation. Data were acquired from three independent experiments and analysed by a one-way ANOVA test. **G.** The proliferation kinetics of control and Vim KO U251 cells after 210 mJ/cm^2^ UV light irradiation was monitored for 7 days by Holomonitor cell proliferation assay. Five positions of two separate wells were acquired per condition. Data were from three independent experiments and analysed by a one-way ANOVA test.

We used then an MTT assay to assess cell viability. This revealed significantly higher survival rates of Vim KO cells compared to control cells following UV exposure (**Fig. 3D**). Additionally, the percentage of cells positive for the apoptosis marker Annexin V was significantly lower in Vim KO cells (**Fig. 3E**), consistent with reduced apoptosis and enhanced cell viability. Ki67 staining and monitoring of cell population growth over seven days post-UV exposure further confirmed this trend: while UV-induced DNA damage effectively halted the expansion of control cells, it only partially reduced the proliferation of Vim KO cells (**Fig. 3F, G**).

Together, these findings demonstrate that vimentin is critical for the induction of DDR pathways, including DNA damage-induced inhibition of cell proliferation and induction of apoptosis. Loss of vimentin allows GBM cells to bypass DNA damage checkpoints, thus contributing to enhanced cell survival and proliferation under conditions of genotoxic stress.

### Vimentin mechanically protects the cell nucleus from compression

The well-established crucial mechanical role of vimentin IFs (Infante and Etienne-Manneville, 2022) and the significant impact of vimentin depletion on gene expression detailed in this study (**Fig. 1**) led us to hypothesize that cytoplasmic vimentin IFs might directly influence nuclear organization and function. In U251-MG GBM cells, vimentin filaments form a distinct perinuclear cage structure (**Fig. 4A**), consistent with observations in other cell types (Patteson et al., 2019). Despite no significant changes in the nuclear volume between control and Vim KO cells (**Supplementary Figure Fig. 3A**), the nuclear morphology was strongly affected. In Vim KO cells plated on glass coverslips, nuclei appeared rounder (as indicated by a decreased aspect ratio) and significantly flatter (reduced height) compared to control cells (**Fig. 4B-E**).

**Figure 4.**
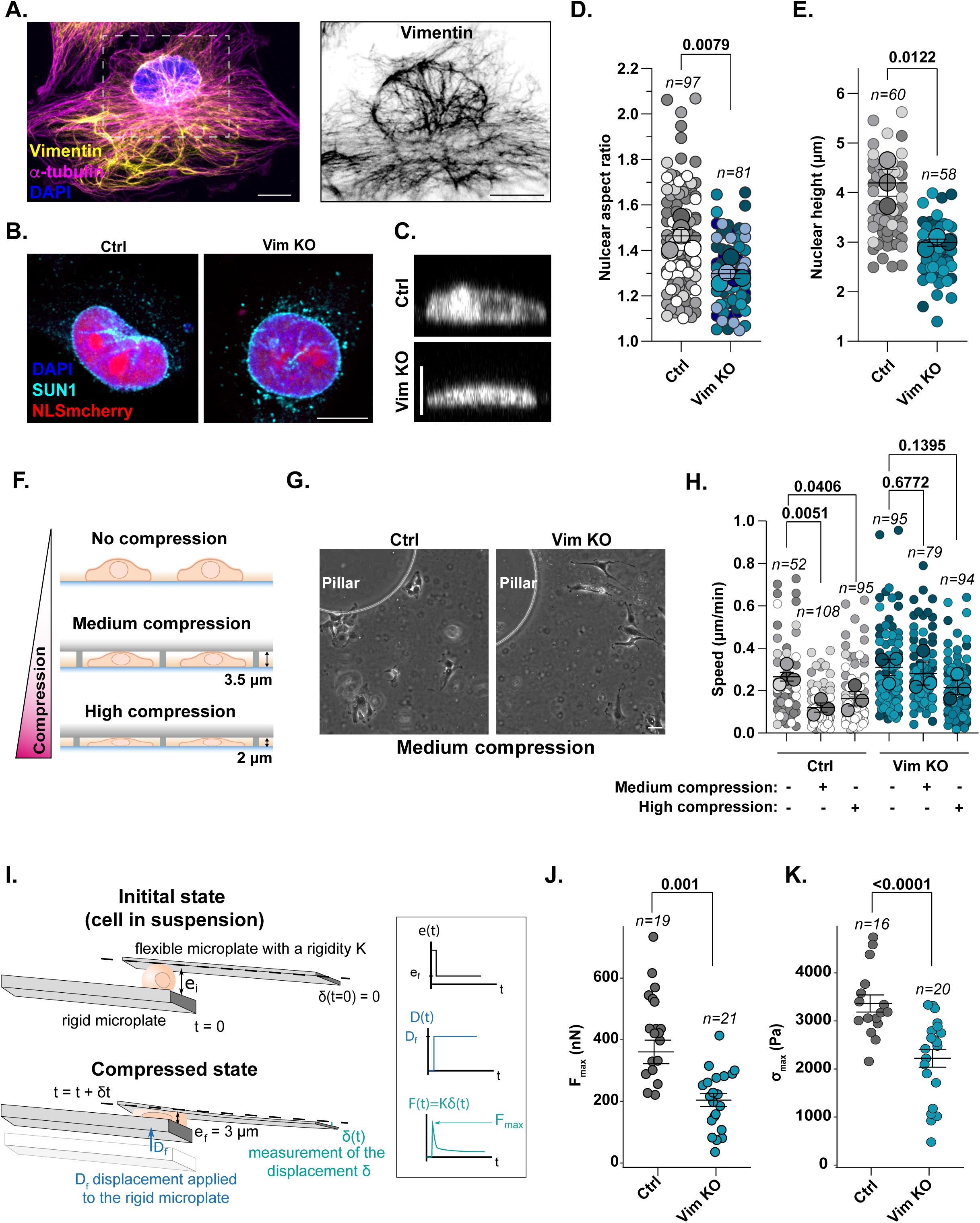
Vimentin perinuclear cage mechanically protects against compression. **A.** (Left) Representative maximum intensity projection of confocal z-stacks of U251-MG cells plated on glass coverslips and stained for vimentin, a-tubulin and DAPI. (Right) Inset of inverted LUT of boxed region showing vimentin staining around the nucleus (scale bar 10 µm). **B.** Representative maximum intensity projection of confocal images of NLS-mCherry control or Vim KO U251-MG cells plated on glass coverslips and stained for SUN1 and DAPI (scale bar 10 µm). **C.** Representative orthogonal view or z-stacks acquired by confocal microscopy of control and Vim KO U251-MG cells. **D and E.** SuperPlots of (D) nuclear aspect ratio and (E) nuclear height. Single data points are represented as small dots in grey (Ctrl) or blue (Vim KO). The mean of each repeat is symbolized by bigger dots. Data were acquired from (D) four or (E) three independent experiments and analysed using a two-tailed *t*-test. *n*=total number of cells per condition**. F.** Representative scheme of the cell confiner device. **G.** Representative images of control and vimentin-depleted U251-MG cells undergoing medium compression (scale bar 50 µm). **H.** SuperPlots of migration speed of control or Vim KO under no, medium (∼3.5 µm) or high (∼2 µm) compression (**Supplementary Video 1 and 2**). Single data points are represented as small dots while the mean of each repeat is symbolized by bigger dots. Data were acquired from three independent experiments and analysed using a one-way ANOVA test. *n*=total number of cells per condition. **I.** Schematic of the microplate rheology used to deform the cell in suspension from its original size to 3 µm thickness and measure the **(J)** maximal force and **(K)** stress needed for this deformation. Data were acquired from three independent experiments and analysed using a two-tailed t-test. *n*=number of cells per condition.

We confirmed the nuclear flattening of vim KO cells by assessing their migratory behaviour under compression. We subjected cells to mechanical compression using a static cell confiner (Le Berre et al., 2014). Cells were confined between two glass coverslips separated by either 3.5 µm or 2 µm, generating “medium” or “high” compression, respectively (**Fig. 4F-H**). In control cells, both medium and high compression substantially reduced migration speed (**Fig. 4H and Supplementary Video 1**). In contrast, the migration speed of Vim KO cells was not significantly affected under either compression conditions (**Fig. 4H and Supplementary Video 2**). Interestingly, compressed Vim KO cells migrated at a rate comparable to non-compressed control cells, suggesting that the loss of vimentin promotes cell migration in a confined compressive environment by facilitating nuclear deformation.

To verify that the impact of vimentin loss on nuclear morphology was due to a change in cell mechanical properties, we used single-cell microplate rheology (Alibert et al., 2021). In this system, individual cells were trapped between two microplates and compressed from their original suspension state (∼10–15 µm thickness) to a final compressed state of 3 µm thickness, to measure the force required to generate such deformation (**Fig. 4I**). Both the maximum force and stress-strain values required to achieve this deformation were markedly lower in Vim KO cells compared to control cells (**Fig. 4J and K**, respectively).

Our findings underscore the critical role of vimentin in mechanically shielding the nucleus from compressive forces. They suggest that vimentin may regulate gene expression by limiting nuclear deformability and mechanotransduction in response to compressive forces.

### Cell compression mimics the effects of vimentin loss on DDR pathways

Mechanical stress is known to alter chromatin organization (Jain et al., 2013; Le et al., 2016). We thus investigated the effects of cell compression (3.5 µm height) on histone modifications using the cell confiner system. Post-translational modifications of histone 3 (H3), including lysine 27 and lysine 9 tri-methylation (H3K27me3 and H3K9me3) and lysine 9 acetylation (AcH3K9), were markedly reduced upon compression of control cells (**Fig. 5A**). Interestingly, vimentin depletion caused a similar reduction in these histone marks, effectively mimicking the effects of mechanical compression (**Fig. 5A**). Immunofluorescence analysis of vim KO cells confirmed the results of the western blot experiments (**Fig. 5B**). Re-expressing human vimentin in Vim KO cells was sufficient to rescue H3K27me3 expression to a similar extent than control condition (**Supplementary Figure Fig. 4A**). Notably, compressing Vim KO cells did not further decrease these histone markers, suggesting that loss of vimentin and nuclear compression potentially act through the same mechanism rather than exerting additive effects. These results suggest that compressive stress phenocopies the loss of the vimentin filaments, leading to significant chromatin modifications.

**Figure 5.**
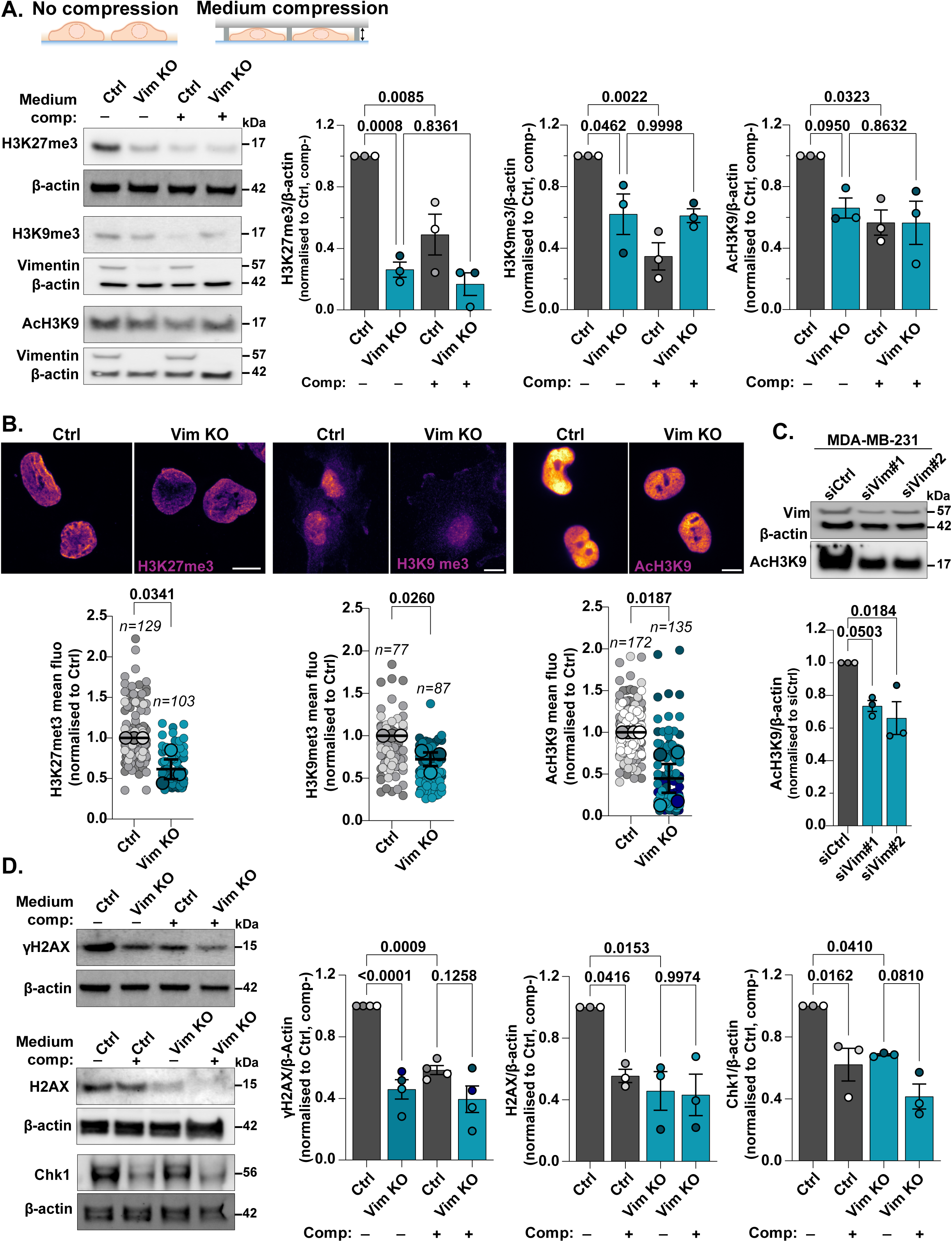
Compression mimics Vimentin depletion of DDR. **A.** (Top) representative scheme of the cell confiner device. (Bottom) Expression of H3K27me3, H3K9me3 and AcH3K9 proteins in control and Vim KO U251-MG cells undergoing or not 4h of medium compression and analysed by western blot. Relative expression was quantified using Fiji. Data were acquired from three independent experiments and analysed using a one-way ANOVA test. **B.** (Left) Representative maximum intensity projection of confocal z-stacks of U251-MG cells plated on glass coverslips and stained for H3K27me3 and representative epifluorescence images (centre and right) of H3K9me3 and AcH3K9 in control and Vim KO U251-MG with relative mean intensity quantification. Data are represented as SuperPlot. Single data points are represented as small dots in grey (Ctrl) or blue (Vim KO). The mean of each repeat is symbolized by bigger dots. Data were acquired from three independent experiments and four independent experiments for AcH3K9 and analysed using a two-tailed *t*-test. **C.** Expression of vimentin and AcH3K9 in MDA-MB-231 cells depleted or not for vimentin using siRNA technology was analysed by western blot analysis and relative expression was quantified using Fiji. Data were acquired from three independent experiments and analysed using a two-tailed *t*-test. **D.** Expression of γH2AX, H2AX and Chk1 proteins in control and Vim KO U251-MG cells undergoing or not 16h of medium compression and analysed by western blot. Relative expression was quantified using Fiji. Data were acquired from four independent experiments and analysed using a one-way ANOVA test.

A similar downregulation of AcH3K9 was observed in vimentin-depleted MDA-MB-231 cells like vimentin loss, exerts a comparable effect on histone marks across different cellular contexts (**Fig. 5C**).

We next evaluated whether mechanical compression of the nucleus could also mimic the effects of vimentin depletion on DDR pathways. Compression of control cells using the static cell confiner led to downregulation of γH2AX, H2AX, and Chk1 protein levels, mirroring the effects observed in Vim KO cells (**Fig. 5D**). To further confirm the impact of compressive stress, we compared DDR protein expression in U251-MG cells grown as three-dimensional (3D) multicellular spheroids versus isolated cells. ATM, Chk1, and H2AX protein levels were significantly lower in spheroids than in isolated cells (**Supplementary Figure Fig. 4B and C**). These results show that vimentin depletion and mechanical compression act through a shared mechanism to impact chromatin organization and DDR pathways.

### Cell compression and vimentin depletion both promote cell survival following DNA damage

Given the strong influence of vimentin and cell compression on DDR pathways, we investigated how vimentin expression and cell compression impact GBM cell survival under various stress conditions encountered during tumour progression.

First, we explored the effects of mechanical stresses encountered during invasion through confined environments, by monitoring cell migration through a polydimethylsiloxane (PDMS) microfluidic chip composed of an array of pillars with a 2 µm inter-pillar spacing (**Fig. 6A**). Vim KO GBM cells traversed constrictions more rapidly than control cells, confirming their increased deformability and enhanced migratory capacity (**Fig. 6B and Supplementary Video 3 and 4**). However, the absence of vimentin increased frequency of nuclear envelope ruptures during migration (**Fig. 6C, 6D and Supplementary Video 5 and 6**). Transient cytoplasmic bursts of a nuclear fluorescent reporter (NLS-mCherry) were twice as frequent in Vim KO cells as in control cells, indicating increased nuclear envelope instability. Interestingly, these experiments also revealed that while the absence of vimentin increases nuclear susceptibility to envelope rupture and potential DNA damage, it does not affect their proliferation **(Supplementary Figure Fig. 5A)** and overall survival (**Fig. 6E**) under mechanical stress.

**Figure 6.**
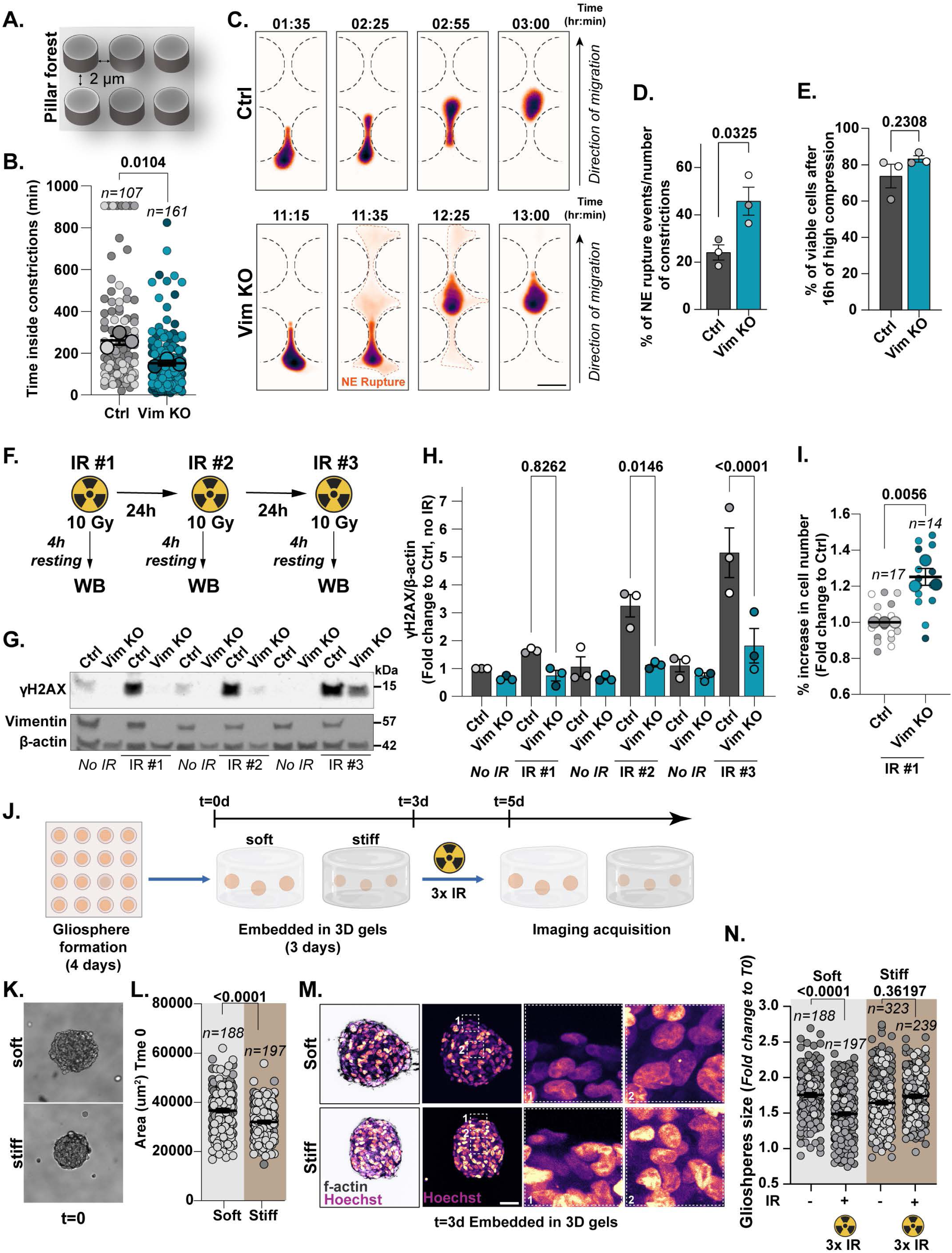
Vimentin downregulation and compression control cell survival. **A.** Schematic representation of the pillar forest microfluidic device. **B.** SuperPlots of the transition time of control and Vim KO cells through 2 µm spaced pillars (**Supplementary Videos 4-5**). Single data points are represented as small dots in grey (Ctrl) or blue (Vim KO). The mean of each repeat is symbolized by bigger dots. Data were acquired from three independent experiments and analysed using a two-tailed t-test. *n*=number of cells per condition. **C.** The gallery shows representative non-consecutive frames from movies obtained from three independent experiments of NLS-mCherry control and NLS-mCherry Vimentin KO cells passing through 2 µm spaced pillars (**Supplementary Videos 5-6**). **D.** Quantification of the nuclear envelope rupture events in control and Vim KO cells during passage through 2 µm spaced pillars. Data were acquired from three independent experiments and analysed using a two-tailed t-test. **E.** MTT assay of control and Vim KO cells after 16h of high compression (2 µm). Data were acquired from three independent experiments and analysed by a two-tailed t-test. **F.** Schematic representation of different doses of X-Ray irradiation (IR). **G-H**. U251-MG control or Vim KO cells were subjected to different doses of X-Ray as indicated, lysed and the protein level of γH2AX was analysed by western blot and quantified using Fiji. Data were acquired from three independent experiments and analysed by a one-way ANOVA test. **I.** Proliferation of cells was quantified after 72h of 1 dose of X-Ray irradiation. SuperPlots represent the % increase in cell number. Single data points are fields of view of Ctrl (grey) or Vim KO (blue). The mean of each repeat is symbolized by bigger dots. Data were acquired from three independent experiments and analysed by a two-tailed t-test. *n*=number of fields of view per condition. **J**. Experimental flow chart of gliosphere irradiation. **K.** Representative bright field images of gliospheres embedded in soft or stiff gel at t=0. **L.** Quantification of gliosphere area. Data were acquired from three independent experiments and analysed by a two-tailed t-test. **M.** Representative maximum intensity projection of confocal z-stacks of gliosphere embedded for 3 days in soft or still gel. Insets shows zoom of selected area. **N.** Quantification of gliosphere size after 3 doses of X-Ray irradiation. Data were acquired from three independent experiments and analysed by mixed linear model.

Standard GBM treatment typically involves surgical resection, radiotherapy, and temozolomide chemotherapy (Stupp et al., 2005; Wen and Kesari, 2008). However, the effects of these interventions are often transient, with recurrence and resistance posing major challenges. This resistance is thought to be driven by the profound intratumoural heterogeneity characteristic of GBM (Bhargav et al., 2021; Pichol-Thievend et al., 2024; Qazi et al., 2017). We thus tested the impact of vimentin expression in the cellular response to repeated X-ray exposures (10 Gy per day over three days) (**Fig. 6F**). γH2AX levels progressively accumulated in control cells following each irradiation, as expected. In contrast, γH2AX levels remained strikingly low in Vim KO cells (**Fig. 6G,H**). Moreover, Vim KO cells exhibited a higher proliferation rate than control cells following irradiation (**Fig. 6I**). This finding suggests that vimentin-negative cells have impaired DNA damage sensing, allowing them to bypass cell cycle checkpoints and evade apoptosis, ultimately enhancing their survival after irradiation.

To determine whether mechanical compression had a similar effect on cell survival following irradiation as vimentin depletion we used GBM cell gliospheres embedded in non-degradable gels of different rigidity (**Fig. 6J**). In ‘stiff’ hydrogels (∼ 3 kPa), gliospheres became more compact than when embedded in ‘soft’ hydrogels (∼ 50 Pa) (**Fig. 6K-M**). After three days (**Fig. 6M**), embedded gliospheres were submitted to 10Gy-irradiations every 24h to mimic therapeutic protocols. Consecutive irradiations significantly reduced gliosphere size in ‘soft’ gels, reflecting substantial cell death, whereas size of gliospheres embedded in ‘stiff’ gels remained unaffected (**Fig. 6N**). These observations strongly suggest that cell compression, which like vimentin loss represses DDR pathways, enhances cell survival following irradiation. Our results reveal that the loss of vimentin in both GBM cells and breast adenocarcinoma cells induces nuclear compression, leading to alterations in histone modifications. This nuclear compression, whether caused by vimentin loss or external compressive forces, significantly suppresses DNA damage response (DDR) pathways and DNA damage checkpoints, thereby enabling sustained cell proliferation and survival under stress conditions (**Fig. 7**). These findings highlight a common mechanism by which the absence of vimentin in the core of the tumour or increased compressive forces allow tumour cells to retain their proliferative capacity despite accumulating DNA damage. This mechanism likely contributes to genomic instability, intratumoral heterogeneity, and resistance to treatment.

**Figure 7.**
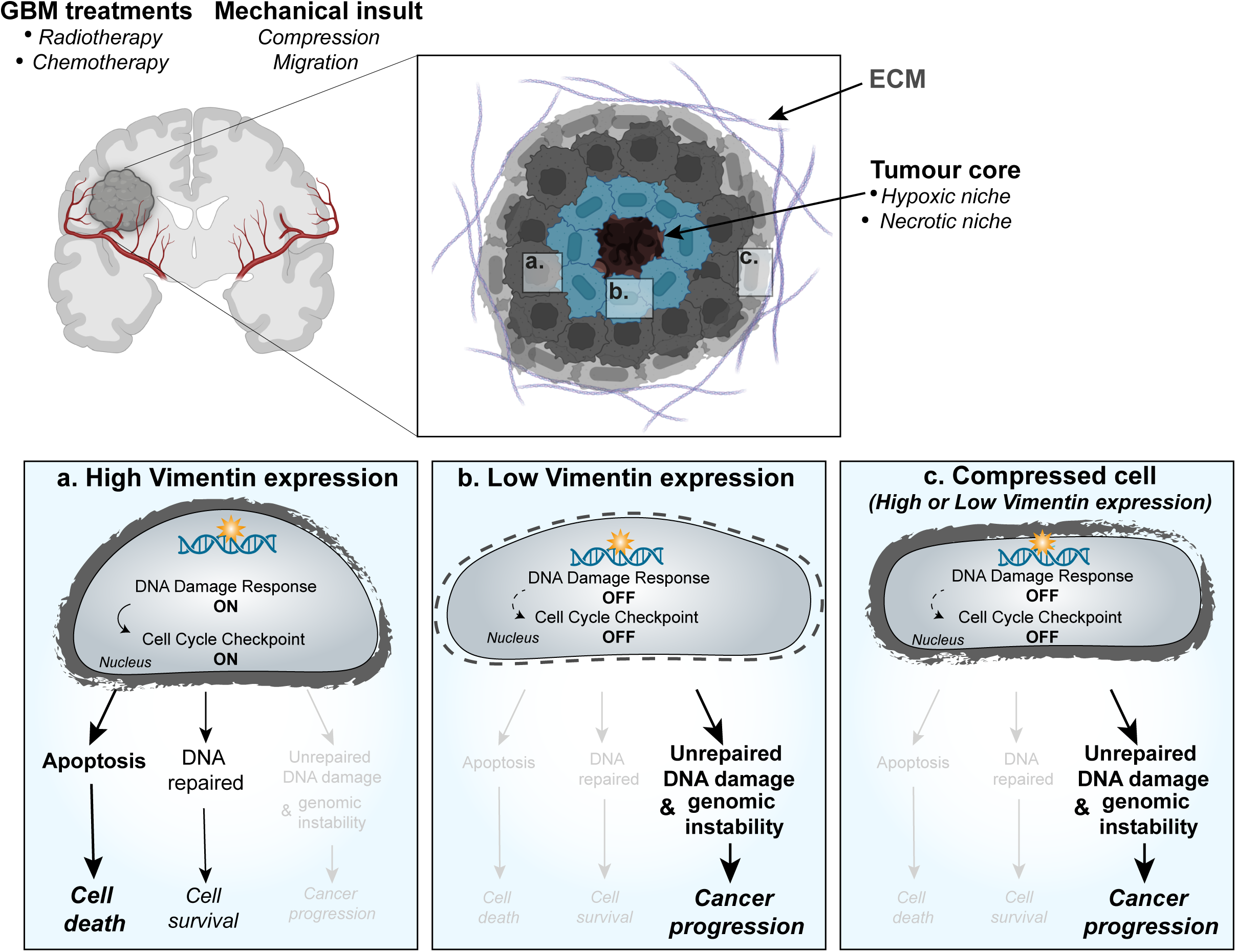
Proposed working model for the role of vimentin during GBM progression. The core of the GBM is considered soft, highly necrotic with vaso-occlusion and hypoxia, while the periphery/ pseudolopalisade and infiltrating rim is characterised by a high elastic module, and ECM secretion. The low expression of vimentin in the core, with a low level of DNA damage sensing machinery and a dysregulation of the cell cycle checkpoint, favours genomic instability and cancer progression in this necrotic and hypoxic environment. Compression at the edge of the tumour, which downregulates the DNA damage sensing machinery similarly to the absence of vimentin favours cell survival.

## Discussion

In this study, we demonstrate that both external compressive forces and the loss of vimentin IFs profoundly impact nuclear morphology, chromatin organization, and DNA damage response (DDR) pathways, ultimately enhancing cell survival under physical stresses, including mechanical compression and X-ray irradiation. The ability of mechanical compression to recapitulate the effects of vimentin loss on histone modifications and on the expression of DDR-related genes strongly indicates that the mechanical properties of the vimentin-based IF network are the primary drivers of these changes.

Previous studies, including our own, have shown that vimentin depletion inhibits the expression of other glial IF proteins (GFAP, nestin, synemin), effectively eliminating the IF network from the cytoplasm(van Bodegraven EJ et al., 2023). The absence of vimentin has been associated with increased cellular deformability in various cell types, including normal fibroblasts and cancerous GBM, and breast adenocarcinoma cells, in which it has been linked to nuclear dysmorphia (Pan and Chen, 2024). Consistent with these findings, our results show that vimentin loss significantly reduces the force required to induce large nuclear deformations under compression (**Fig.4I,4K**). This effect likely arises from the role of the perinuclear vimentin IF cage (**Fig. 4A**), which acts to limit the transmission of mechanical forces from the cytoplasm to the nucleus (Pan and Chen, 2024; van Bodegraven EJ et al., 2023). Additionally, the contribution of vimentin IFs to the overall elasticity of the cytoplasm may also participate to their buffering capacity (Infante and Etienne-Manneville, 2022). The similar and non-additive effects of vimentin depletion and high compressive forces on histone modifications and DDR gene expression, strongly suggest that the vimentin filaments act purely mechanically. However, we cannot fully exclude that part of vimentin-dependent transcriptomic regulation may involve biochemical pathways as described for keratin17 (Hobbs et al., 2015), or the direct effect of vimentin as a transcription factor (Fan et al., 2023).

Compressive forces exerted on the cell nucleus, whether through external mechanical stress or the loss of vimentin-mediated resistance, induce nuclear deformation and regulate the epigenetic state through mechanotransduction-dependent pathways (Lv et al., 2018). Epigenetic regulation governs chromatin structure, influencing the balance between transcriptionally inactive heterochromatin and transcriptionally active euchromatin regions (Hu et al., 2022; Uhler and Shivashankar, 2017). These chromatin states are dynamically modulated, and their dysregulation can significantly affect cellular processes such as differentiation, proliferation, and responses to environmental stress or damage (Hu et al., 2022). However, the precise mechanisms by which nuclear compression influences chromatin remodelling enzymes and transcriptional machinery, thereby the resulting gene expression patterns, remain to be elucidated.

Our main result is the fact that nuclear compression, whether it is induced by the loss of vimentin IFs or by external compressive forces or by both, leads to a dramatic downregulation of all major DDR pathways. These pathways include key DNA damage sensors and signalling proteins such as ATM, which activate downstream effectors to halt the cell cycle and facilitate repair or trigger apoptosis if the damage is irreparable. In addition to the downregulation of ATM, we observe a strong inhibition of H2AX. H2AX phosphorylation plays a pivotal role in the DNA damage response, particularly in the repair of double-strand breaks (DSBs). Its phosphorylation at serine 139 by ATM, ATR, or DNA-PK to form γH2AX acts as a molecular beacon to amplify the DDR signal, recruiting and retaining DNA repair proteins at the damage site. Here, not only are the major DDR kinases downregulated but H2AX expression is also strongly reduced, leading to a dramatic decrease in γH2AX, even when DNA damage is strongly induced by UV or X-Rays (**Fig. 3 or 6**, respectively). In addition to the alteration of DNA damage sensing, DNA repair proteins, including 53BP1, are also downregulated. Decreased expression and activation of these tumour suppressors, which normally ensure the stability of the genome, likely leads to accumulation of genomic alterations (Gorodetska et al., 2019; Gudmundsdottir and Ashworth, 2006). Finally, PARP1, which is directly involved in the repair of single-strand DNA breaks to prevent these lesions from converting into more dangerous double-strand breaks during DNA replication, is also downregulated, likely also contributing to genomic instability (Ray Chaudhuri and Nussenzweig, 2017). Our results show that GBM cells that lack vimentin or are submitted to large compressive forces lack most of the DNA damage sensing pathways and thus cannot efficiently repair DNA, leading to persistent DNA lesions. Accumulation of such damage increases genomic instability, which can drive tumour progression and promote the emergence of cancer subclones with adaptive advantages.

Defects in double strand DNA break repair should activate cell cycle checkpoints or apoptosis pathways as a protective mechanism. However, when exposed to UV, X-Rays or constraints-induced nuclear envelop leakage, Vim KO cells proliferate more and undergo less apoptosis than control cells. This surprising observation can be explained by the downregulation of major targets of the DDR pathway which should mediate cell cycle checkpoints. For instance, loss of vimentin decreases the serine/threonine kinase Chk1 (**Fig 1 and 2**), which phosphorylates and inhibits CDC25 to induce cell cycle arrest (Uto et al., 2004). Moreover, 84% of GBM patients and 94% of GBM cell lines, including U251-MG cells display altered p53-ARF-MDM2 pathway (Zhang et al., 2018) and low expression of vimentin has been correlated with a decrease in p53 (Sembritzki et al., 2002), suggesting that the p53 pathway is generally inactivated in vimentin negative GBM cells. Finally, apoptosis is also diminished in Vim KO cells. Although we did not detect any major changes in the level of expression of Bcl2 family members, there was an inhibition of APAF-1, another major player of the apoptotic intrinsic pathway (**Fig. 1E**). Inhibition of APAF-1 not only prevents procaspase-9 activation and apoptosis, but it also prevents mitochondrial damage, which may also contribute to the cell survival that we observed following UV radiations **(Fig. 3E)** (Gortat et al., 2015). We propose that the profound alteration of the DDR pathways, together with a downregulation of the downstream signalling normally leading to cell cycle arrest or cell death leads to the accumulation of DNA damages while the cell keeps proliferating. Ultimately, this will likely result in some level of cell death, but it may also strongly contribute to intratumoral heterogeneity.

In GBM, cancer cells are softer than healthy astrocytes. Notably, grade IV GBM cells, the most invasive, are softer than grade III astrocytoma cells (Alibert et al., 2021). While these cells are softer overall due to reduced cytoskeletal protein levels, they exhibit increased perinuclear stiffness due to the local accumulation of actin and IFs. The lower expression of vimentin within the tumour core compared to the periphery (**Fig. 1B**) suggests that this region may be prone to the accumulation of mutations, whereas the higher proportion of vimentin-positive cells at the tumour periphery would promote invasion. However, we previously showed that vimentin-positive cells could lead the migration of vimentin-negative cells (Pan and Chen, 2024; van Bodegraven EJ et al., 2023). This suggests that vimentin-negative cells may also disseminate and contribute to tumour recurrence.

Importantly, cell compression can trigger similar effects as the loss of vimentin. We did not detect any changes in vimentin expression level following overnight compression, but we cannot exclude than in the long term, such regulatory mechanisms may also be at play. As GBM grows within the brain tissue, it exerts a local pressure on its surroundings and results in a tissue displacement known as the gross mass effect, responsible for the acute neurological symptoms seen in patients. However, because the space occupied by the brain is restricted by the cranium, tumour growth also implies compression of the tissue (Ropper, 1986). We show that gliospheres growing within a stiffer environment (thus, a more limited space) have better resistance against X-Ray exposure **(Fig. 6N)**, suggesting that this mechanical compression could also promote tumour radioresistance.

Since loss of vimentin also lead to the alteration of nuclear shape, of chromatin marks and to the downregulation of DDR pathways in GBM and breast adenocarcinoma cells (**Supplementary Figure Fig. 2 and Fig. 5C)**, we propose that the mechanisms described here may be at play in several cancer types (Pan and Chen, 2024). Altogether these findings underscore the pivotal role of cancer mechanics in disease progression. Both the mechanical properties of the tumour microenvironment and of the tumour cells themselves significantly influence tumour cell resistance to DNA damage, probably contributing to radiotherapy resistance and promoting tumour heterogeneity. These mechanical factors also likely impact tumour responses to chemotherapies targeting DNA damage response (DDR) pathways, as softer or compressed cells can evade apoptosis and continue proliferating despite impaired DNA repair mechanisms.

Moreover, our results highlight vimentin as a key player in tumour progression. While vimentin expression facilitates tumour cell invasion, its absence paradoxically enhances tumour cell resistance to therapy and promotes genomic alterations. These findings suggest that vimentin expression could serve as a valuable molecular marker to differentiate tumour cells with distinct behaviours, potentially guiding the selection of the most effective treatment strategies. This work paves the way for further exploration of vimentin as both a diagnostic tool and a therapeutic target in cancer management.

## Acknowledgements

We gratefully acknowledge the UtechS Photonic BioImaging (Imagopole), C2RT, Institut Pasteur, supported by the French National Research Agency (France BioImaging; ANR-10–INBS–04; Investments for the Future). We are very thankful to the Biomics Platform, the Bioinformatics and Biostatistics Hub, Unit of Technology and Service for Cytometry and Biomarkers (UTechS CB), and the Biomaterials and Microfluidic platform of Institut Pasteur, for their contribution to this project. We thank the Human Glioblastoma Cell Culture resource (www.hgcc.se) for the patient-derived U3117 glioblastoma cells. We thank Philippe Chavrier for MDA-MB-231 cells, Pedro Monteiro for comments on the manuscript, Giorgio Seano and the PCMC lab members for insightful discussions.

## Funding

This work was supported by La Ligue contre le cancer (EL2023 - DN/IP/IQ – 17691), Worldwide Cancer Research (WCR 23-0156), INCA PLBIO (PLBIO24-072), INCA PCSI (N° 22CP073-00) and Institut Pasteur (PTR-548-22). EI was supported by Fondation de France (WB-2021-35925) and INCA PCSI (N° 22CP073-00), EvB by NWO VI.Veni.212.117

## Author contribution

Conceptualization: EI, SEM

Methodology: EI, ET, GM, SH, DP, VR, SK, RR, EvB, SEM

Investigation: EI, ET, GM, SH, DP, VR, SK, RR, EvB

Supervision: EI, EvB, AA, SEM

Writing— original draft, review & editing: EI, SEM

## Competing interests

All other authors declare they have no competing interests.

## Ethics, Consent to Participate, and Consent to Publish declarations

Not applicable

## Data availability

All the raw data and code generated during this study are available from the corresponding author on reasonable request.

The single cell RNA sequencing data from Yuan et al., 2018 and Darmanis et al., 2017 are available in the Gene Expression Omnibus under the accession GSE103224 (Yuan et al., 2018) and GSE84465 (Darmanis et al., 2017).

The raw and processed RNA sequencing data of U251 WT and Vim KO cells are available on GEO under accession number GSE286944 (https://www.ncbi.nlm.nih.gov/geo/query/acc.cgi?acc=GSE286944) using the following password: svmvuqmytvsbxyp

## Materials and methods

### Cell culture, stable cell lines and siRNA treatment

U251-MG cells were grown in Minimum Essential Medium (MEM) with GlutaMAX supplement (Thermo Fisher Scientific) supplemented with 10 mM of MEM non-essential amino acid, 100 U ml−1 penicillin, 100 mg ml−1 streptomycin (Gibco) and 10% fetal calf serum (Eurobio Scientific) at 37 °C under 5% CO2. U3117 cells were grown on Geltrex (Gibco) pre-coated plates and grown in serum-free GBM complete medium composed of Neurobasal-A medium mixed 1:1 (v/v) with DMEM/F12 + GlutaMAX and complemented with human FGF-basic and human EGF at 20 ng/mL, and B-27 (Gibco).

Vimentin KO U251-MG cell line was generated as described before (Jiu et al., 2017). Guide RNAs (sgRNAs) to target Vimentin as described in Jiu et al., 2015 with high on-/off-target ratios for different exons were used. The pSpCas9(BB)-2A-GFP plasmid (PX458, Addgene, Catalog #48138) containing a gRNA scaffold under the U6 promoter and Cas9 and EGFP under the CMV promoter was linearized with BbsI-HF (NEB) and specifically designed annealed oligonucleotides containing the guide sequences (sgRNA 1, targeting exon 2: 5’ CACCGTGGACGTAGTCACGTAGCTC-3’ and 5’-AAACGAGCTACGTGACTACGTCCAC-3’, sgRNA 2, targeting exon 2: 5’-CACCGCAACGACAAAGCCCGCGTCG -3’ and 5’-AAACCGACGCGGGCTTTGTCGTTGC -3’) were ligated into the linearized plasmid using T4 Ligase (Thermo Fischer Scientific).

Patient-derived U3117 GBM cells were acquired from the Human Glioblastoma Cell Culture resource (www.hgcc.se) at the Dept. of Immunology, Genetics and Pathology, Uppsala University, Uppsala, Sweden (Xie et al., 2015). Cells were transfected with the sgRNA-containing plasmids by Lipofectamine 2000 (ThermoFisher Scientific) according to the manufacturers’ protocol. Cells were plated in culture dishes and maintained for three days. GFP-positive cells were subsequently collected using fluorescent activated cell sorting (FACS) and plated as single cells in 96-well plates and were grown till confluency and maintained. Vimentin knockout was determined using immunostaining and western blot analysis. sgRNA 2 resulted in the highest number of vimentin KO clonal lines. A control and KO clonal line were selected based on the absence of vimentin protein and an IF-network and were used for analyses.

U251-MG cells stably expressing NLS-mCherry were generated by nucleofection (Amaxa Biosystems Nucleofection System) with 1 µg of DNA in 100 µl of Glial nucleofection reagent (Lonza Biologics), followed by selection with 500 µg ml^−1^ geneticin (Thermo Fisher Scientific). After fluorescence-activated cell sorting (FACS), polyclonal cell lines were obtained and cultured in complete media supplemented with 500 µg ml^−1^ geneticin.

Vimentin-rescued U251-MG cell lines were generated by re-expression of vimentin cDNA in vimentin-depleted cells from Vimentin-7-EGFP (Addgene #56439), modified by introducing two stop codons between the vimentin sequence and the EGFP reporter sequence. Cells were transfected using Lipofectamine 2000 (Thermo Fisher Scientific) according to the manufacturers’ protocol. Three days post-transfection, cells were treated with 500 µg ml^−1^ geneticin to select transfected cells. A single cell clone was generated and selected based on immunoreactivity for vimentin.

MDA-MB-231 were grown in DMEM-High Glu medium supplemented with 100 U ml−1 penicillin, 100 mg ml−1 streptomycin and 10% fetal calf serum at 37 °C under 5% CO2.

U251-MG and MDA-MB-231 cells were transfected by nucleofection using 1nmol siRNA (Sigma-Aldrich) in 100 μl of Nucleofection reagent (kit C, Lonza Biologics, Slough, UK). Experiments were carried out 72h post-transfection. siRNA sequences used in this study are listed as followed Luciferase (siCtrl)-UAAGGCUAUGAAGAGAUAC; siVim#1-UAAGCUCUCUAGUUCUUAACA; siVim#2-UGAAGAAACUCCACGAAGA

### Analysis of GBM patient-derived published single-cell RNA sequencing data

To explore vimentin expression in patient-derived GBM samples, a large integrated GBM single-cell RNA sequencing dataset, GBM Map24 was used. The Seurat object containing integrated and annotated data was downloaded (Ruiz-Moreno, 2022) and used for analyses using Seurat 5.0.1. Samples from recurrent GBM were excluded from analyses. To determine the percentage of vimentin positive and negative cell per patient, raw read counts were used. Cells with > 1 raw count for vimentin were annotated as vimentin positive cells, while cells with < 1 raw count for vimentin were annotated as cells in which vimentin was not detected (negative cells).

### RNA sequencing

RNA from cells was extracted on ice using a RNeasy Mini kit (Qiagen) according to the manufacturer’s instructions and treated with DNase. RNA concentration was determined by measuring the ratio of the absorbances at 260 and 280 nm. Quality control of RNA samples was assessed with a 2100 Bioanalyser system according to manufacturer’s instructions.

Libraries were prepared from 300 ng of RNA using Illumina Stranded mRNA Prep (Illumina) following the manufacturer’s protocol. Briefly, mRNA was captured by oligo-dT magnetic beads, fragmented and reverse transcribed. Sample-specific barcodes were ligated to the cDNA and then amplified by 17 cycles of PCR. Note that only the first cDNA strand was amplified. Purification of unbound adaptors and primers was done on AMP beads (Beckman Coulter) added to samples in a 0,8:1 ratio. The resulting stranded libraries comprised fragments from 200 to 1000 bp with a peak around 490 bp, as visualized on a 5300 Fragment Analyser (Agilent Technologies). Libraries were pooled, purified on AMP beads as described above, diluted to 0,65 nM and sequenced on a NextSeq 2000 system (Illumina) using a P3 50-cycle sequencing kit. The yield was 1100 mln 67-bp single-end reads (30 – 60 mln reads per sample)

### RNA-seq and statistical analysis

The RNA-seq analysis was performed with Sequana 0.14.1 (Cokelaer et al. 2017). We used the RNA-seq pipeline 0.18.1 (https://github.com/sequana/sequana_rnaseq) built on top of Snakemake 7.8.5 (Köster and Rahmann 2012). Briefly, reads were trimmed from adapters using Fastp 0.20.1 (Chen et al., 2018) then mapped to the Homo sapiens GRCh38 genome assembly from Ensembl using STAR 2.7.8a (Dobin et al., 2013). FeatureCounts 2.0.1 (Liao et al., 2014) was used to produce the count matrix, assigning reads to features using corresponding annotation v94 from Ensembl with strand-specificity information.

RNAseq count data were analysed using R version 4.2.1(R, 2024) and the Bioconductor package DESeq2 version 1.36.0 (Love, 2024). The normalization and dispersion estimation were performed with DESeq2 using the default parameters and statistical tests for differential expression were performed applying the independent filtering algorithm. A generalized linear model was set in order to test for the differential expression between VIM and control. Raw p-values were adjusted for multiple testing according to the Benjamini and Hochberg (BH) procedure (Hochberg, 1995) and genes with an adjusted p-value lower than 0.05 were considered differentially expressed. Gene set enrichment analysis was performed using the fgsea R package (Korotkevich G, 2019; Subramanian et al., 2005) on the Wikipathways gene-sets(Agrawal et al., 2024). Only gene sets with a FDR lower than 0.05 were considered significantly enriched.

### Western blot analysis

Cells were lysed using Laemli buffer with protease and phosphatase inhibitor cocktails. Lysates were separated by SDS–PAGE and transferred to a polyvinylidene difluoride membrane. Membranes were incubated in blocking buffer (3% BSA in PBS containing 0.1% Tween-20) for 1 h at room temperature (RT) and then incubated overnight at 4°C with the indicated antibodies. Antibodies were visualized using either the ECL detection system or fluorescent secondary antibodies (Bio-Rad). Signals were recorded using a ChemiDoc MP Imaging System (Biorad). Densitometric analysis was performed using FIJI.

### Immunofluorescence microscopy

Sample were fixed with 4% PFA at RT for 10 min or, for microtubule staining, with cold methanol at -20 °C for 10 min. Samples were then washed three times with PBS 1×. Blocking was performed using a 5% BSA in PBS 1× solution for 1h at RT. The primary antibodies diluted in 5% BSA in PBS 1× were then incubated 1h at RT. After three washes using 5% BSA in PBS 1×, samples were incubated with secondary antibodies for 1h at RT. Samples were then washed three times in PBS 1×1×, and mounted with ProLong Gold antifade reagent with DAPI. Images were acquired either by epifluorescence microscopy using a Leica DM6000 microscope equipped with a 40X 1.25 NA or 63X 1.4 NA objective or using a LSM 700 confocal microscope with a plan-Apochromat 40X/1.4 Oil objective.

### Nuclear volume quantification

Nuclear volume analysis was performed using a custom FIJI macro. The macro processes TIFF images by segmenting the nuclei through thresholding and image binarization, ensuring accurate delineation of nuclear boundaries. After segmentation, 3D reconstruction of the nuclei is performed based on the image stack to account for volumetric data. Their volumes is then calculated based on predefined input parameters, with the results saved in a designated output folder.

### Cell compression

Cells were plated on a 6 well plates a glass bottom dishes (MatTek, P06G-1.5-20-F) 16 h prior confinement using a commercial confiner (4D cell, France). Confiner slides were generated internally. The template for the slides was fabricated by soft photolithography according to the protocol previously described (Heuze et al., 2011). To prepare the slides, a drop of polydimethylsiloxane mixture (PDMS RTV 615 A and cross-linker B at a ratio of 10:1 (w/w), Neyco, France) was placed on the mold that contained the pillars of target height (∼3.5 µm or ∼2 µm) and immediately covered with a 12-mm glass coverslips. The wafers were baked at 95 °C for 15 min and coverslips covered with PDMS pillars were carefully removed using isopropanol. After washing them with water and letting them dry, coverslips covered with PDMS pillars were placed on the soft spacer used to link the lid of the dish to the bottom of the well and incubated in cell medium for 2 h before confinement. Cells were either imaged by time-lapse microscopy or lysed after 4 h or 16 h of compression. Migration speed (μm/min) was obtained by tracking cells using the Manual Tracking Fiji analysis software plugin.

### Pillar forest microfluidic device

Pillar forests were prepared from molds fabricated in the same maner as the ones used in the compression experiments (see Cell compression section in Material and Methods). The microfluidic chip was prepared by casting 5 g of a mixture of PDMS RTV615 (Neyco, France) at a ratio 10:1 w/w PDMS A:crosslinker B. The mixture was then degassed vacuum and cured at 65 °C for 1 h. The chip was then removed from the mold and resized to fit the glass part of a glass-bottom 35 mm dish (Fluorodish FD-035, WPI) and 2.5 mm holes were punched out (Biopunch, TedPella) in order to inject cells in the chip. After sonication of the PDMS chip in 70% ethanol, the glass and the chip were air-plasma treated for 30s at max power (Harrick PDC-002). Both activated surfaces were then put in contact to ensure sticking. To strengthen the bonding, the dish containing the chip was set in an oven at 65 °°C for 1. Finally, the chip was washed with PBS and incubated with complete medium for at least 1 h before adding 100 000 cells in each entry.

### X-Ray and UV irradiation

Cells or spheroids where irradiated using an X-Ray generator (Faxitron) at the indicated doses or with an UV (312 nm) irradiation system (UVITEC, Cambridge).

### Comet assay

Comet assay was performed following manufacture instructions (Abcam, ab238544). Briefly, agarose drops were placed on a comet slide to form a base layer. Subsequently, cells subjected to UV irradiation where mixed with agarose and seeded on top of the base layer. Following polymerisation at 4 °C, cells were lysed and the slides were placed in an electrophoresis tank in alkaline solution. After electrophoresis, samples were stained using a DNA dye and imaged with DMI6000B Leica epifluorescence inverted microscope equipped with10X objective and a Photometrics MOMENT sCMOS camera controlled by Micromanager (Edelstein et al., 2014).

### MTT assay

Cells were plated overnight on a 96-well plate. Cells were exposed to either UV or high compression forces and after 16 h tetrazolium salt was added according to the manufacturer’s instructions (Roche). Optical density was measured at 570 nm with Safas M96 plate reader and analysed with Safas software.

### Live-cell imaging

Time-lapse images were acquired with 10X or 20X objective using a Nikon Eclipse Ti-E epifluorescence inverted microscope with an EMCCD camera/pco.edge sCMOS camera and Metamorph software. Cells were maintained under 5% CO_2_ at 37°C. Images were acquired every 5 or 15 min for 16 h according to the experimental setup. Cells were manually tracked with Fiji software (Manual Tracking plug-in).

### HoloMonitor

1×10^4^ cells were plated on a12 well plate (Sarstedt) and holographic images were acquired for 7 days according to the HoloMonitor cell proliferation protocol. Five positions of two separate wells were acquired per condition. Automated quantification was obtained using the HoloMonitor App Suite.

### Generation of spheroids

Spheroids were prepared using a MicroTissues silicone mold (12-256 - Small Spheroids, MicroTissues). 2% molten agarose was poured into the silicone mold and allowed to cool in the refrigerator for 5 min. The agarose cuvettes were then demolded into a 6 well plate and soaked in PBS 1X supplemented with 1% penicillin-streptomycin and stored at fridge temperatureat 4°C until use. The agarose cuvettes were placed in an incubator before the dissociation of the cells to bring them to 37 °C.

The PBS 1X solution was aspirated just before depositing the cell solution. The spheroids were generated by pipetting 150 µL of dissociated cell suspensions in agarose cuvettes at a concentration of 106 cells/mL. Then, complete GBM medium was added and the spheroids were left to grow for 4 days under 5% CO_2_ at 37 °C. After 4 days, the generated spheroids were gently flushed out of the cuvette with medium and collected in a falcon tube. They were then centrifuged at 1000 RPM for 3 min to pellet them down. The supernatant was gently removed and 200 µL of fresh medium was then added to resuspend the spheroids.

### Spheroids encapsulation in hydrogel

The hydrogel solution was prepared following the protocol provided by Cellendes. Two types of hydrogel were generated: the “soft” hydrogel, using the fast-gelling dextran hydrogel at a final concentration of 1.5 mM (FG90-1, Cellendes GmbH), and the “stiff” hydrogel, using the slow-gelling dextran hydrogel at a final concentration of 3 mM (G92-1, Cellendes GmbH). The solutions were left to polymerize at 37 °C for 10 or 15 min, respectively, before the addition of fresh medium. The encapsulated spheroids were then left to grow for 2 days before the first series of irradiation.

### Statistical analysis

Graphs were generated using GraphPad Prism (10.2.1). Statistical tests used in this study is indicated in each figure legend.

## Notes

### Competing Interest Statement

The authors have declared no competing interest.

