## Supplementary Figures for "Intratumoral Heterogeneity of Vimentin Modulates Nuclear Mechanotransduction, DNA Damage Response and Cancer Cell Survival"

### Supplementary figure legends.

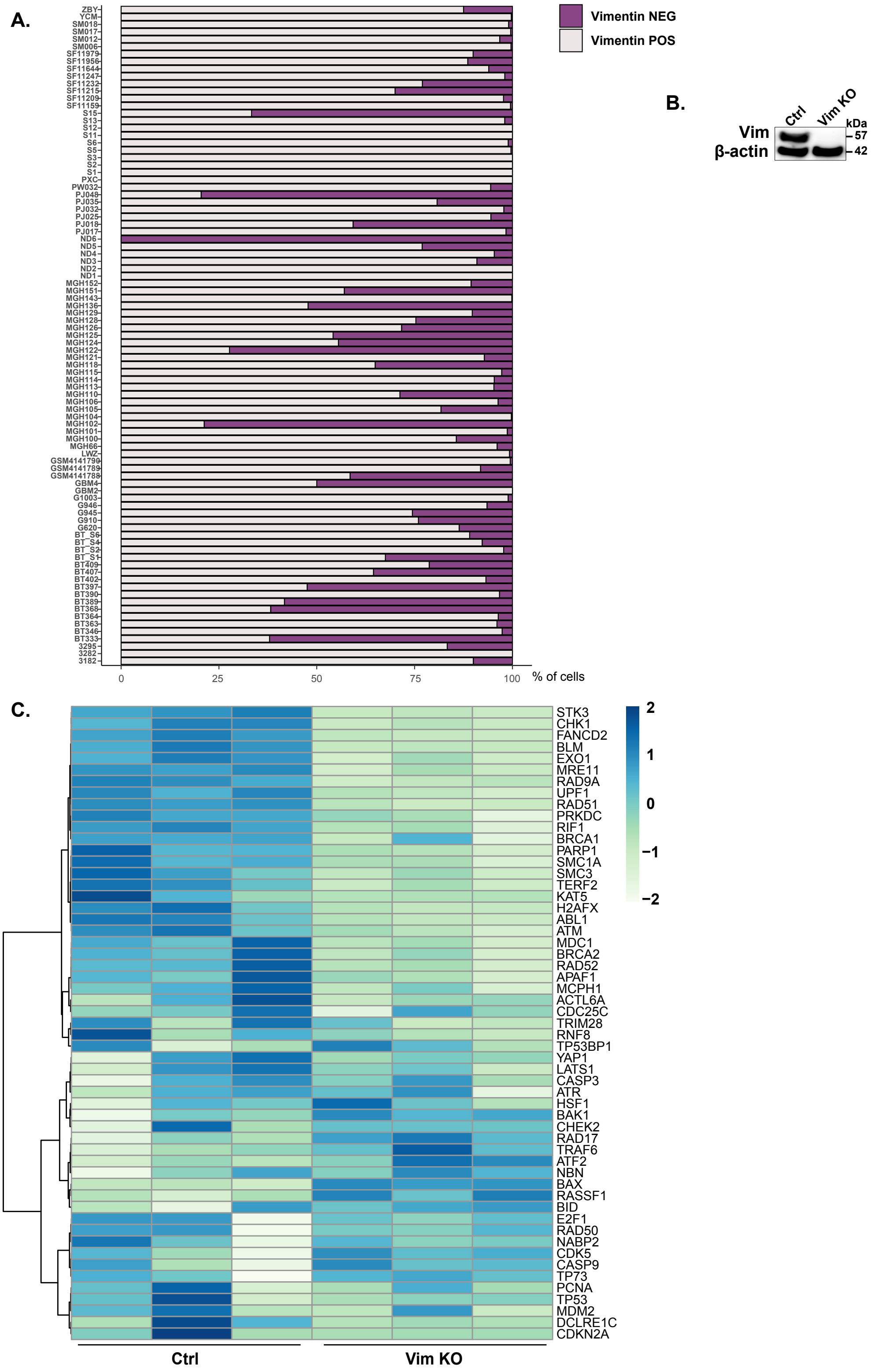


**Supplementary figure 1. A.** Percentage of vimentin-positive cells per patients from single cell RNA sequencing dataset. **B.** Expression of vimentin protein in control and Vim KO U251-MG cells was analysed by western blot. **C.** Heatmaps providing a full visual representation of gene expression of the DNA IR-double strand breaks and cellular response via ATM pathways in control versus Vim KO cells.

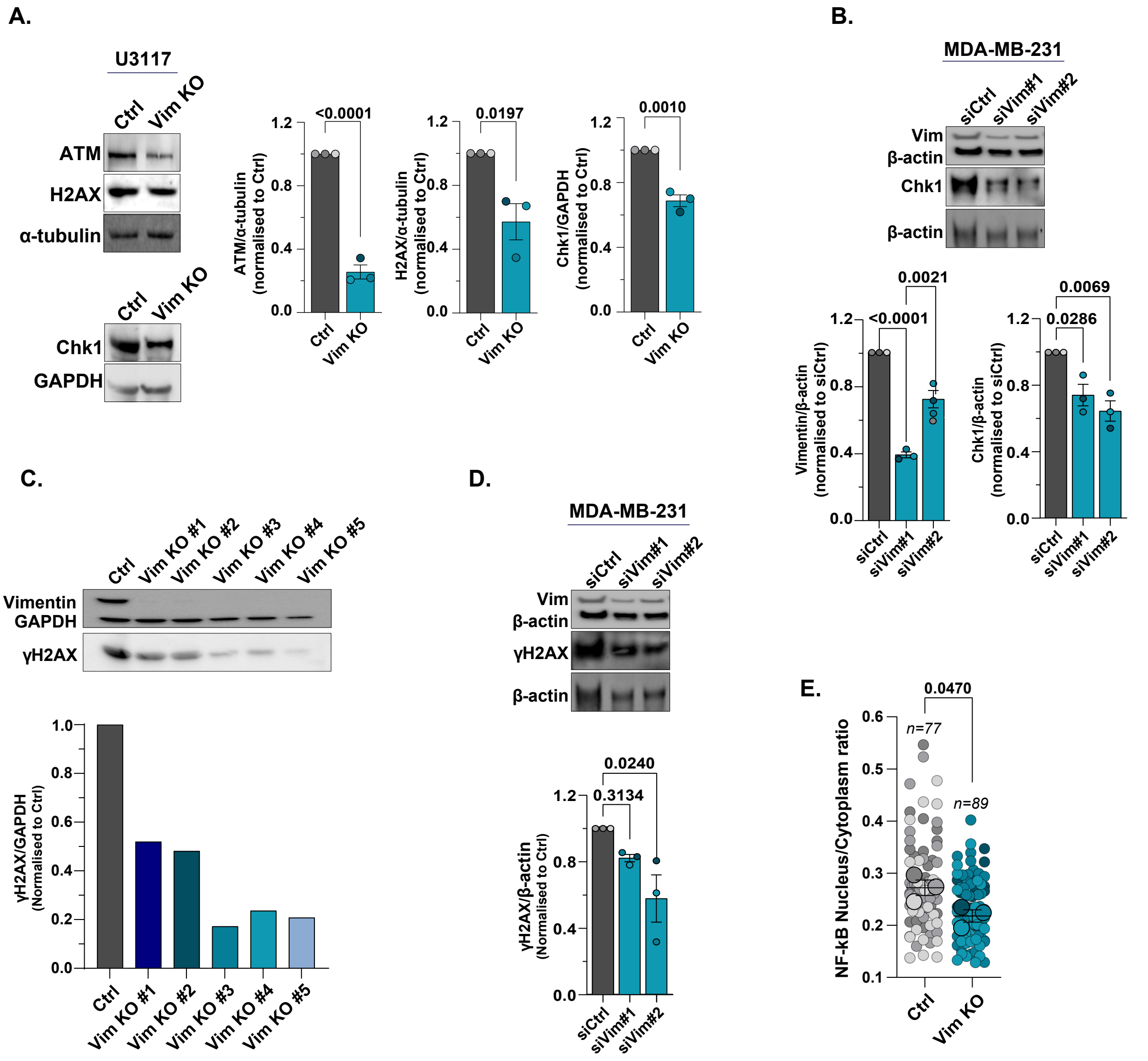


**Supplementary figure 2. A.** Expression of ATM, H2AX and Chk1 in control and Vim KO U251-MG cells was analysed by western blot and relative expression was quantified using Fiji. Data were acquired from three independent experiments and analysed using a two-tailed *t*-test. **B.** Expression of Chk1 in MDA-MB-231 cells depleted or not for vimentin using siRNA technology was analysed by western blot and relative expression was quantified using Fiji. Data were acquired from three independent experiments and analysed using a two-tailed *t*-test. **C.** Expression of γH2AX in control and five single clones of Vim KO U251-MG cells was analysed by western blot and relative expression quantified using Fiji. **D.** Expression of γH2AX in MDA-MB-231 cells depleted or not for vimentin using siRNA technology was analysed by western blot and relative expression was quantified using Fiji. Data were acquired from three independent experiments and analysed by a one-way ANOVA test. **E.** SuperPlots of nucleus versus cytoplasm ratio of NF-kB measured by immunofluorescence analysis. Single data points are represented as small dots in grey (Ctrl) or blue (Vim KO). The mean of each repeat is symbolized by bigger dots. Data were acquired from three independent experiments and analysed by a two-tailed *t*-test. *n*=total number of cells per condition.

**A.**

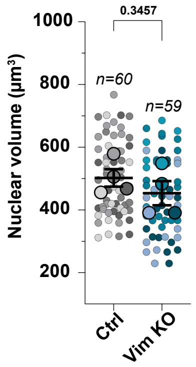


**Supplementary figure 3. A.** SuperPlots of nuclear volume in control and Vim KO of U251-MG cells. Single data points are represented as small dots in grey (Ctrl) or blue (Vim KO). The mean of each repeat is symbolized by bigger dots. Data were acquired from four independent experiments and analysed by two-tailed *t*-test. *n*=total number of cells per condition.

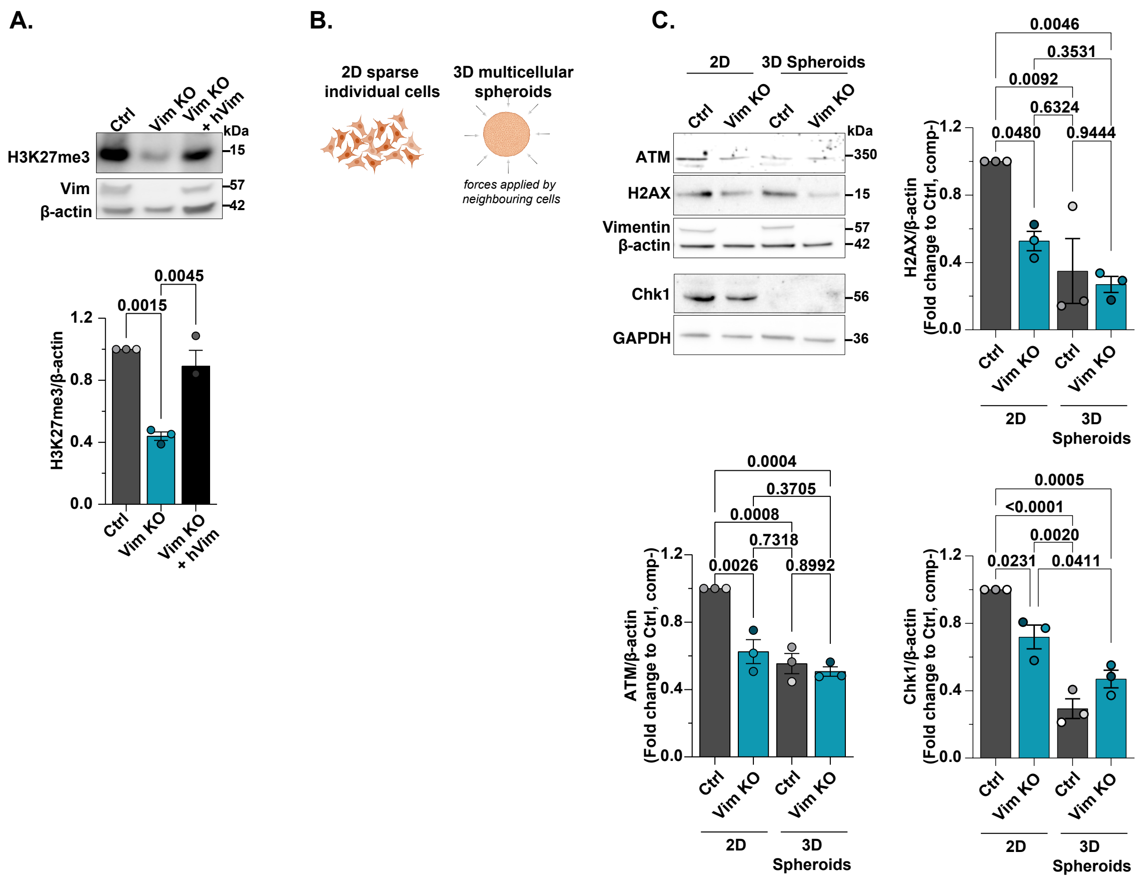


**Supplementary figure 4. A.** Expression of H3K27me3 protein in control, Vim KO and vimentin rescue in U251-MG cells analysed by western blot. Relative expression was quantified using Fiji. Data were acquired from three independent experiments and analysed using a one-way ANOVA test. **B.** Schematic representation of experimental design comparing sparse 2D cell culture to 3D spheroids culture. **C.** Expression of ATM, Chk1 and H2AX in control and Vim KO U251-MG cells cultured as sparse 2D cells or as spheroids was analysed by western blot and relative expression was quantified using Fiji. Data were acquired from three independent experiments and analysed using a two-tailed *t*-test.

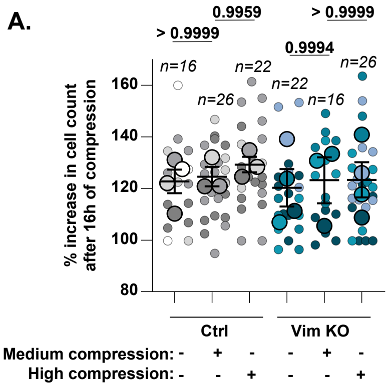


**Supplementary figure 5. A.** Proliferation of cells was quantified 16h of no, medium or high compression. SuperPlots represent the % increase in cell number. Single data points are fields of view of Ctrl (grey) or Vim KO (blue). The mean of each repeat is symbolized by bigger dots. Data were acquired from three or four independent experiments and analysed using a one-way ANOVA test. *n*=number of fields of view per condition.

**Antibodies list.**

| Antibodies | Source | Reference |
|---|---|---|
| Mouse monoclonal anti-Vimentin | Santa Cruz Biotechnology | sc-6260 |
| Mouse monoclonal anti-β-actin | Sigma-Aldrich | A2228 |
| Mouse monoclonal anti-GAPDH | Merck | MAB374 |
| Rat monoclonal anti-a-Tubulin | Biorad | MCA77G |
| Rabbit polyclonal anti-H2AX | Cell signaling | 2595 |
| Rabbit polyclonal anti-pH2AX (S139) | Cell signaling | 2577 |
| Mouse monoclonal anti-pH2AX | Merck | JBW301 |
| Rabbit polyclonal anti-ki67 | Abcam | 15580 |
| Rabbit monoclonal anti-ATM | Cell signaling | 2873 |
| Mouse monoclonal anti-lamin A/C | Santa Cruz Biotechnology | 376248 |
| Rabbit monoclonal anti-H3me3K27 | Cell signaling | 9733 |
| Rabbit monoclonal Acetyl-Histone H3 (Lys9) | Cell signaling | 9643 |
| Rabbit monoclonal anti-H3me3K9 | Cell signaling | 13969 |
| Mouse monoclonal anti-PCNA | Cell signaling | 2586 |
| Rabbit monoclonal anti- NF-kB | Cell signaling | 8242 |
| Rabbit polyclonal anti-SUN1 | Sigma-Aldrich | HPA008461 |
| Mouse monoclonal anti-Chk1 | Cell signaling | 2360 |
| Rabbit polyclonal anti-Chk1 | Bethyl laboratory | 300-298A |
| HRP- Donkey Anti-rabbit | Jackson ImmunoResearch | 711-035-152 |
| HRP- Donkey Anti-Mouse | Jackson ImmunoResearch | 715-035-150 |
| Tetramethylrhodamine (TRITC) donkey anti-rabbit | | (711-025-152) |
| Alexa Fluor 488 donkey anti-mouse | Jackson ImmunoResearch | 715-545-151 |
| Anti-actin hFAB Rhodamine antibody | Bio-Rad | 12004163 |
| Alexa Fluor 647 phalloidin | Thermo Fisher Scientific | A22287 |
| Annexin V-FITC | Abcam | Ab14085 |
